# Noncanonical loops regulated by EMF1 and cohesin-associated factors shape the distinct 3D genome architecture in plants

**DOI:** 10.64898/2026.02.08.704707

**Authors:** Dingyue Wang, Suxin Xiao, Lingxiao Luo, Jinqi Wei, Jiayue Shu, Guangmei Tian, Yali Liu, Minqi Yang, Tingting Yang, Myriam Calonje, Hang He, Yuannian Jiao, Yue Zhou

## Abstract

In mammals, cohesin-mediated chromatin loop extrusion is blocked at boundaries, where CTCF, WAPL, and PDS5s co-bind and regulate cohesin, establishing an impermeable barrier with no exceptions. In contrast, our study revealed that in Arabidopsis, some noncanonical contacts have emerged between the boundary flanking regions, which suggest that cohesin is not absolutely blocked but enables sliding beyond boundaries. We identified these interactions as noncanonical loops and clarified their regulatory proteins.

Firstly, the plant-specific boundary binding protein EMF1 is essential for noncanonical loops formation. Surprisingly, although both Arabidopsis WAPLs and PDS5s inhibit loop extension as mammals, PDS5s do not co-bind and cooperate with WAPLs at boundaries. Instead, PDS5s bind to boundary-flanking regions enriched with H3K4me1 through their Tudor domains, restricting further extension of noncanonical loops. Consequently, *pds5a/b/c* mutants lose noncanonical loops, resulting in emergence of larger canonical loops but anchored on boundaries and co-upregulation of anchored gene pairs. Collectively, our findings reveal that the plant-specific noncanonical loop, regulated by plant-specific EMF1 and PDS5 proteins, is the fundamental structure of the distinctive chromatin organization of plants, which provides novel mechanistic insights into eukaryotic chromatin architecture regulation.

## Introduction

The cohesin complex was first identified as an essential component of sister chromatid cohesion, a critical process for chromosome segregation during cell division^1,2^. Subsequent research revealed that cohesin also plays roles in organizing the three-dimensional (3D) chromatin structure^1,3,4^, DNA damage repair^5^, and transcriptional regulation^6^. The cohesin complex consists of four conserved components: structural maintenance of chromosomes 1 (SMC1), SMC3, sister chromatid cohesion 1 (SCC1, encoded by the α-kleisin gene), and SCC3^7,8^. SMC1 and SMC3 function as ATPases and form a V-shaped heterodimer. SCC1 and SCC3 complete this structure, forming the cohesin ring^1,7,9^. In mammals, cohesin mediates chromatin loop formation through loop extrusion, a process typically halted at boundaries via interactions with CCCTC-binding factor (CTCF)^10,11^. This boundary-paused cohesin is further regulated by boundary-bound factors Wings Apart-Like (WAPL) and Precocious Dissociation of Sisters 5 (PDS5)^12^. While WAPL promotes cohesin release^13^, PDS5 exhibits complex functionality: it restricts loop elongation through facilitating release or by inhibiting cohesin’s enzymatic activity^14–16^. Collectively, the cooperation of CTCF, WAPL, and PDS5 ensures that cohesin-mediated loop extrusion does not cross boundaries, a principle widely observed across all studied eukaryotes^11,12^. Although cohesin components and their extrusion mechanism are conserved across eukaryotes^17,18^, the regulatory mechanisms governing cohesin-dependent loops in plants remain largely unknown.

In *Arabidopsis*, the cohesin components SMC1, SMC3, and SCC3 are encoded by single-copy genes, whereas α-kleisins are represented by four homologs (SYNAPTIC 1-4, SYN1-4)^19^. Mutations in *SMC1*, *SMC3*, and *SCC3* result in lethal phenotypes, indicating that cohesin plays a vital role^20–23^. The *syn1* mutant is sterile but shows minimal developmental abnormalities, while mutations in *SYN3* cause gametophyte lethality^24–28^. SYN2 and SYN4 display functional redundancy in DNA double-strand break repair^29^; *SYN4* exhibits the highest stable transcript levels among all α-kleisin genes during mitosis^29,30^ and directly binds a DNA G-box motif^31^. Unlike mammals, *Arabidopsis* lacks CTCF homologs. Instead, EMBRYONIC FLOWER1 (EMF1) acts as a genome architectural protein that interacts with cohesin to maintain 3D chromatin structures^32^. Furthermore, both cohesin-associated factors, WAPLs and PDS5s, are functionally conserved in chromatin organization regulation^33^. *Arabidopsis* possesses two *WAPL* genes, and the *wapl1/2* mutant exhibits only mild developmental and reproductive defects^34^. Five *Arabidopsis* functionally redundant *PDS5* genes are essential for genome organization: Hi-C analyses reveal global chromatin reorganization in *pds5* mutants, including aberrantly enhanced TAD-like structures^33^. Notably, *Arabidopsis* PDS5C contains a Tudor domain with demonstrated H3K4me1-binding capacity^35^, providing a mechanistic basis for its chromatin regulatory role.

Studying the 3D genome of *Arabidopsis* has been challenging due to its distinct chromatin organization compared to mammals^36,37^. The *Arabidopsis* genome exhibits mild segregation and predominantly forms local interactions^32,38–40^, making TAD identification particularly difficult. This unique organization and its regulatory mechanisms remain poorly understood. In this study, we specifically identify noncanonical loops as key determinants of *Arabidopsis’* distinctive 3D genome architecture.

## Results

### Noncanonical loop is a novel type of cohesin-dependent loop

The enhanced 3C-based method Micro-C identifies chromatin loops at a resolution of ∼200 bp, exceeding the ∼1 kb resolution of Hi-C and thereby providing more detailed information^41^. Consistent with this enhanced resolution, chromatin loops anchored by SYN4 that were identified using Micro-C could not be resolved by Hi-C analysis (Fig.1a). The high quality and reproducibility of our Micro-C libraries were validated by calculating the number of valid pairs and stratum-adjusted correlation coefficient (Extended Data Fig.2a,b). To investigate the role of cohesin in mediating chromatin loops in *Arabidopsis*, we used the *syn4* mutant as a loss-of-cohesin function model (Extended Data Fig.1), considering the lethality of *smc1*, *smc3*, and *scc3* mutants^20–23^. Interaction decay exponent (IDE) curves showed a significant reduction in contact frequency at genomic distances less than 100 kb in the *syn4* mutant (Fig.1b). Therefore, we defined chromatin loops anchored by SYN4 and spanning less than 100 kb as cohesin-dependent loops in *Arabidopsis*.

**Fig. 1:**
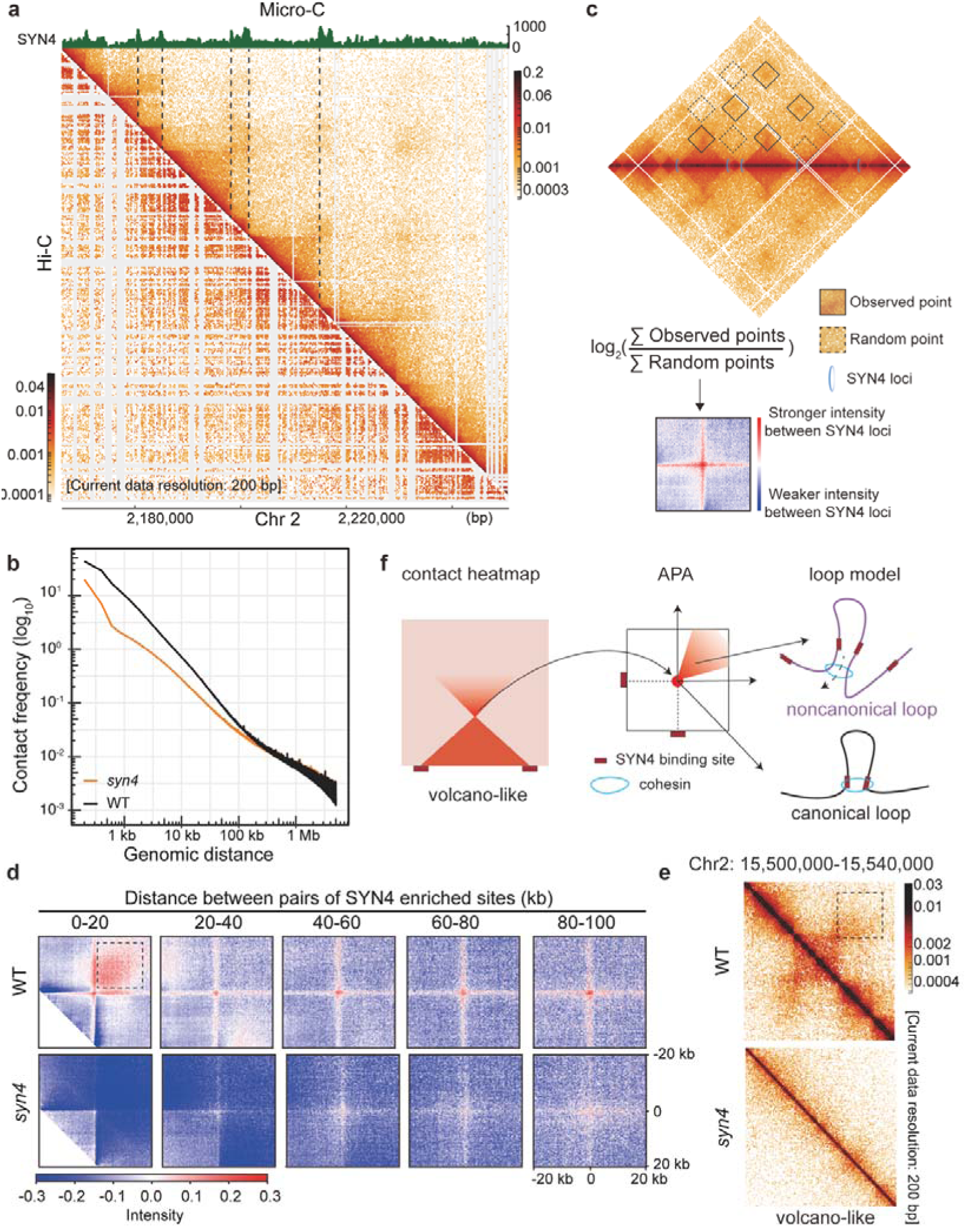
Cohesin-dependent loops include canonical and noncanonical loop in Arabidopsis. a, At a resolution of 200 bp, Micro-C data (top-right heatmap) revealed loop structures forming between SYN4 binding boundaries (dash lines). These loops are not discernible at the same resolution in Hi-C data (bottom-left heatmap). **b,** Decay of contact probability as a function of genomic distance, calculated for the WT (black) and *syn4* mutant (yellow). **c,** Schematic illustrated the APA calculation process. Pairwise interactions between SYN4 boundaries within a given distance range were extracted as observed interactions; interactions between random sites within the same range were used as the random background. Observed interactions were normalized against the random background to generate the APA heatmap. The APA plot was calculated at 1-kb resolution with a 20-kb flanking region. Red indicates stronger interactions between SYN4 binding sites, whereas blue represents interactions occurring more frequently in the random background. **d,** APA plots for the WT and *syn4* mutant at different distances, calculated at 1-kb resolution with a 20-kb flanking region around all SYN4 anchors. Dotted rectangle outlined the signals of noncanonical loops. **e,** Heatmaps showed canonical loops (triangle at the bottom) and noncanonical loops (above the triangle and outlined by the dotted rectangle) in WT; and both loops were eliminated in *syn4* mutant. **f,** Schematic diagrams illustrated the contact heatmap (corresponding to panel 1e) and the APA plot (corresponding to panel 1d). Arrows indicated how these signals align with the two loop models. Black and purple lines represent the canonical and noncanonical loop; red squares denote SYN4 anchors, blue circles indicate the cohesin complex.

To better characterize these loops and statistically assess their correspondence with cohesin, we used SYN4 binding sites in the wild type (WT) as the boundary and performed aggregate peak analysis (APA) at various distances (0-100 kb) (Fig.1c,d). In APA analyses, we established the plot center as the origin of a rectangular coordinate system and delineated quadrants for each corresponding heatmap (Fig.1d,f). WT plants exhibited a distinct central signal in APA plots corresponding to contacts between SYN4 binding sites, manifesting as a triangular structure in the heatmaps (Fig.1e,f). We define these boundary-anchored loops (corresponding to the central signal in APA / triangular vertex in heatmap) as canonical loops, similar to mammalian loops that cannot cross boundaries. Importantly, these canonical loops were completely absent at all genomic distances in the *syn4* mutant (Fig.1d,e), confirming their cohesin dependency. Additionally, distinct loop-like signals (outlined with the dotted rectangle), corresponding to the surrounding chromatin architecture in quadrant I (Fig.1d) and contact signals above the triangle (Fig.1e), were specifically detected at distances less than 20 kb (Fig.1d). These loops together with the below triangular structure and form a novel volcano-like pattern on the heatmaps (Fig.1e,f). Crucially, they cross the SYN4-binding boundary and were also absent in the *syn4* mutant, further supporting their cohesin dependence (Fig. 1d,e). Notably, such noncanonical loops have not been reported in mammals or other eukaryotes. As a result, we suspected the noncanonical loop is the fundamental structures of the distinct chromatin structure in plants. To verify the hypothesis, we further investigated the mechanisms enabling boundary crossing and restricting loop extension of noncanonical loops.

### EMF1 is essential for the formation of noncanonical loops

In mammals, CTCF prevents cohesin-mediated loop extrusion from crossing boundaries by binding to these boundary elements^42^. Although *Arabidopsis* lacks a CTCF homolog, previous studies have shown that plant-specific modulator EMF1 interacts with cohesin and binds to the boundaries^32^. To further characterize the role of EMF1 in noncanonical loop formation, we analyzed cohesin-dependent loops using Micro-C in the *emf1* mutant (Extended Data Fig.1,2a and 2c). IDE curves for the *emf1* mutant also exhibited significantly reduced contact frequencies at genomic distances less than 100 kb, though to a lesser extent than observed in the *syn4* mutant (Fig.2a). Moreover, cohesin-dependent loops, including both canonical and noncanonical loops, were significantly reduced in the *emf1* mutant (Fig.2b,c).

**Fig. 2:**
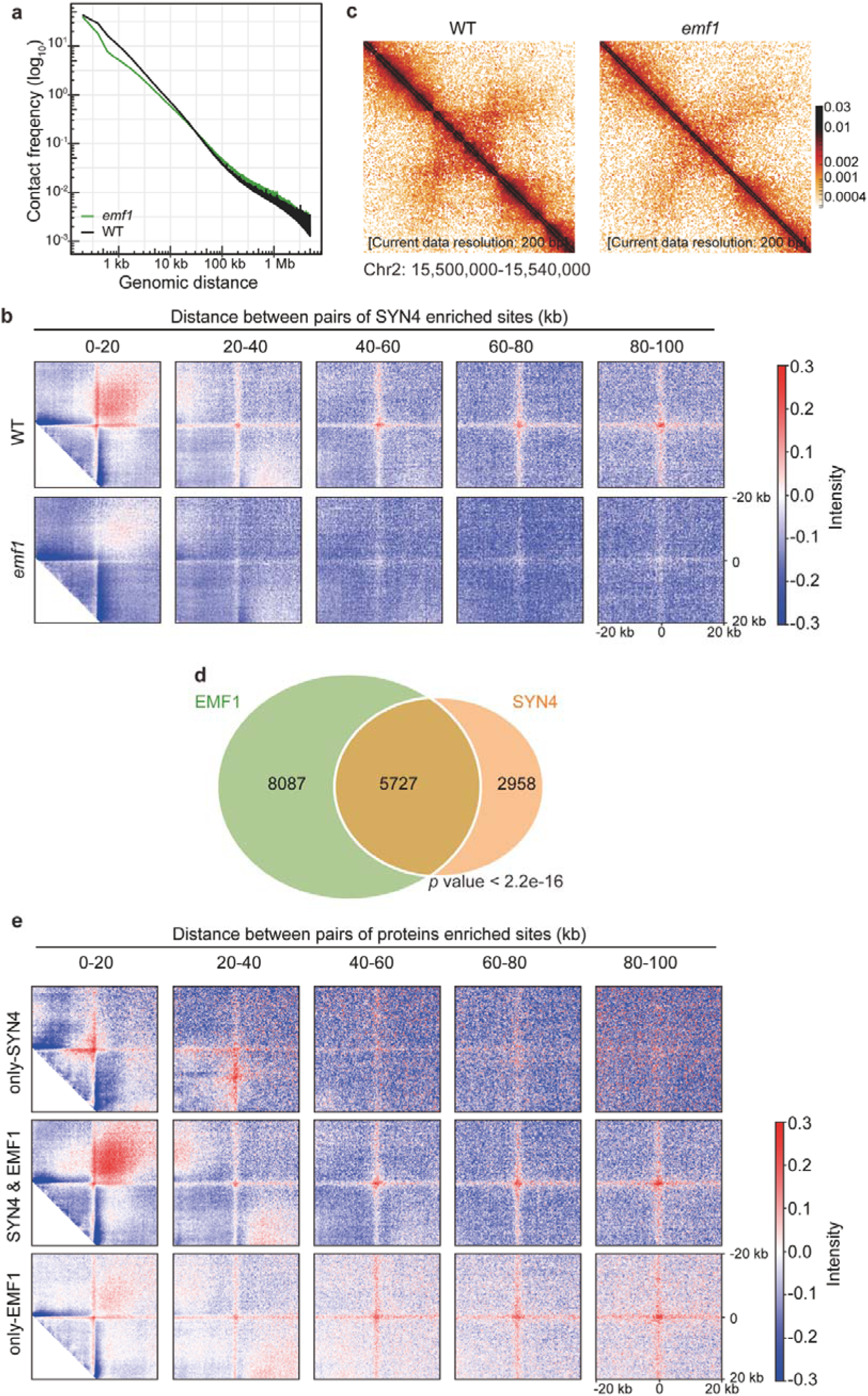
EMF1 takes important role in noncanonical loop formation. **a**, Decay of contact probability as a function of distance, calculated for the WT and *emf1* mutant. **b,** APA analysis in the WT and *emf1* mutant at various distances, calculated at 1-kb resolution with 20-kb flanking regions, focusing on all SYN4 anchors. **c,** Heatmaps showed a reduction in both canonical loops and noncanonical loops in the *emf1* mutant compared to WT. **d,** Venn diagram depicted the overlap between EMF1 and SYN4 binding sites. Statistical significance was calculated using a hypergeometric test. **e,** APA analysis in the WT at 1-kb resolution with 20-kb flanking regions. Panels show APA plots for: anchors bound only by SYN4 (2958 sites, top), anchors co-bound by SYN4 and EMF1 (5727 sites, middle), and anchors bound only by EMF1 (8087 sites, bottom).

Given that the binding sites of EMF1 and cohesin do not completely overlap^32^, we sought to elucidate the mechanisms by which EMF1 regulates cohesin-dependent loops. For this purpose, we compared cohesin (SYN4) and EMF1 peak distributions. We found that approximately 66% of SYN4 peaks (5727 of 8685) overlapped with EMF1 peaks, whereas 34% (2958 of 8685) were SYN4-specific (Fig.2d). We also identified EMF1-only binding sites (8087 of 13814 peaks). Next, we performed APA on sites bound by SYN4 alone, by both EMF1 and SYN4, and by EMF1 alone (Fig.2e). Surprisingly, noncanonical loops were detected only when anchors were co-bound by both SYN4 and EMF1 (Fig.2e). This finding, combined with the reduced noncanonical loop signal observed in the *emf1* mutant (Fig.2b,c), demonstrates that EMF1 is essential for enabling cohesin-mediated loop extrusion cross boundaries to form noncanonical loop. Furthermore, stronger and longer canonical loops were formed between co-bound peaks compared with only-SYN4 bound peaks, indicating that EMF1 contributes to loop expansion (Fig.2e). Loop signals between sites bound only by EMF1 were weak (Fig.2e), and these signals were absent in the *syn4* mutant, confirming their dependence on cohesin (Extended Data Fig.3). Taken together, these results demonstrate that EMF1 functions on cohesin-dependent loop extension and is required for the formation of noncanonical loops.

**Fig. 3:**
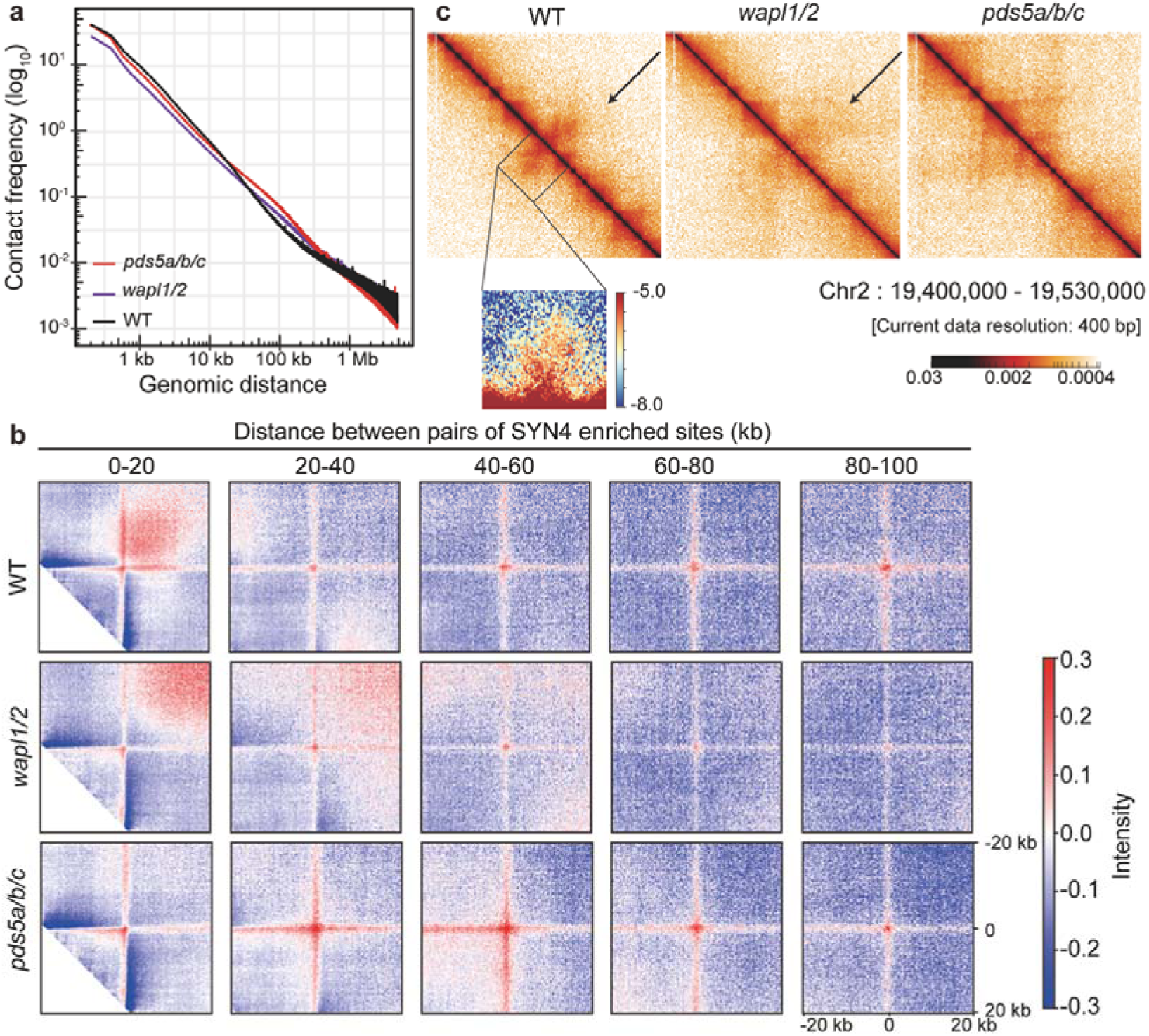
WAPL1/2 and PDS5s restrict noncanonical loop extension through distinct mechanisms. a, Distance-dependent decay of contact probability calculated for the WT, *wapl1/2* and *pds5a/b/c* mutants. **b,** APA for the WT, *wapl1/2* and *pds5a/b/c* mutants at different distances, calculated at 1-kb resolution with 20-kb flanking regions around all SYN4 anchors. **c,** Chromatin interaction heatmaps comparing loop structures: WT (400 bp resolution on the top, 200 bp resolution at the bottom): canonical loops and noncanonical loops are present; *wapl1/2* mutant (400 bp resolution): weakened canonical loops and lengthened noncanonical loops (indicated by the black arrow); *pds5a/b/c* mutant (400 bp resolution): enlarged canonical loops (manifested as expanded triangular structures) with loss of noncanonical loops.

### PDS5s restrict noncanonical loops specifically

In mammals, cohesin-mediated loop extrusion is constrained by the coordinated action of WAPL and PDS5s, which bind to boundary sites to terminate loop extension and catalyze cohesin release at these sites^12,14,15,43^. To further dissect the role of WAPLs and PDS5s on noncanonical loop formation in *Arabidopsis*, we performed Micro-C analysis in the *wapl1/2* and *pds5s* mutants (Extended Data Fig.1,2a,2d and 2e). In mammals, WAPL is involved in releasing cohesin from chromatin at loop anchors^43^. Consistent with this function, IDE curves showed reduced contacts at short distances and increased long-range contacts (Fig.3a). Similarly, APA plots revealed weaker center signals and extended first-quadrant signals towards the upper-right corner; longer noncanonical loops became visible when using larger flanking regions in the APA analysis in the *wapl1/2* mutant (Fig.3b and Extended Data Fig.4). Heatmaps confirmed that chromatin loops in the *wapl1/2* mutant were weaker at short distances but stronger at longer distances compared to the WT (Fig.3c). These results suggest that loss of WAPLs allows a subset of canonical loops to extrude further, forming noncanonical loops; however, these loops become constrained from further extrusion upon crossing boundaries and do not reach subsequent boundaries. Although *WAPL* mutation universally extends cohesin-dependent loops in both *Arabidopsis* and mammals, a fundamental divergence emerges: mammalian loops remain anchored to subsequent boundaries, whereas those in *Arabidopsis* predominantly extend across boundaries yet fail to reach the subsequent boundaries, suggesting the existence of a plant-specific mechanism constraining noncanonical loops again.

In mammals, PDS5s restricts loop elongation by binding to boundaries in addition to WAPL-mediated loop extension restriction^12,43^. To investigate constraints limiting further extension of noncanonical loops to adjacent boundaries in plants and characterize their anchor properties, we analyzed Micro-C datasets from the *pds5s* mutant (Extended Data Fig.1,2a and 2e). IDE curves revealed significantly increased long-range contacts (Fig.3a). APA plots further revealed loss of noncanonical loops, while canonical loops not only remained stable but exhibited strengthened intensity and extended span in the mutant (Fig.3b). Heatmaps showed substantial structural alterations in *pds5a/b/c* mutants compared to WT, manifesting as enlarged triangular structures (Fig.3c). These results indicate that PDS5 proteins act as noncanonical loop extension restriction factors in *Arabidopsis*, with their mutation causing noncanonical loops to extrude toward subsequent boundaries (Fig.3b,c). Collectively, although both WAPL1/2 and PDS5 proteins restrict chromatin loop extension, they regulate distinct loop types: WAPL1/2 regulate canonical loops, preventing the formation of additional noncanonical loops; whereas PDS5s specifically restrict noncanonical loops.

### Distinct occupancy of WAPLs and PDS5s contributes to their different loop extension restriction functions

Given that both WAPLs and PDS5s function as loop extension restriction factors yet exert distinct effects on cohesin-dependent loops, we investigated their binding patterns in relation to loop formation dynamics. For this purpose, we generated genome-wide WAPL1 binding sites through ChIP-seq analysis (Extended Data Fig.5a) and integrated these with published PDS5C ChIP-seq datasets^35^. Feature distribution analysis revealed fundamental divergence in genomic localization: WAPL1 exhibited preferential enrichment at transcription start site (TSS) regions, whereas PDS5C showed pronounced association with gene exons (Extended Data Fig.5b,c).

We next compared their binding patterns relative to SYN4 binding sites. WAPL1 demonstrated substantial overlap with SYN4 loci (Fig.4a,b), a conserved feature in plants and mammals, facilitating canonical loop release. Conversely, PDS5C displayed distinct localization, enriching at flanking regions rather than directly at SYN4 binding boundaries (Fig.4a,c). To characterize PDS5-mediated regulation of noncanonical loops, we categorized all cohesin-dependent loops into four clusters based on PDS5C and EMF1 binding relative to SYN4 binding sites. Notably, Cluster 1, containing the highest number of SYN4 sites (5013) bound by EMF1 and flanked by PDS5C, formed noncanonical loop (Fig.4d). In contrast, SYN4 binding sites lacking EMF1 co-binding failed to establish noncanonical loops despite the presence of PDS5C at flanking regions (Cluster 2, 1645) (Fig.4d), reaffirming EMF1’s indispensable role. Strikingly, Clusters 3 (436) and 4 (1069), where PDS5C was bound directly to SYN4 sites (with or without EMF1), exhibited complete absence of noncanonical loops (Fig.4d). These findings establish that noncanonical loops require two non-redundant conditions: (1) EMF1 is essential for noncanonical loop formation (2) PDS5 restricts noncanonical loop extension at boundary flanking regions. This positions PDS5-enriched regions as functional anchors distinct from boundary-anchored canonical loops, thereby resolving the core mechanistic questions of: (i) boundary crossing drivers and (ii) extension limitation.

**Fig. 4:**
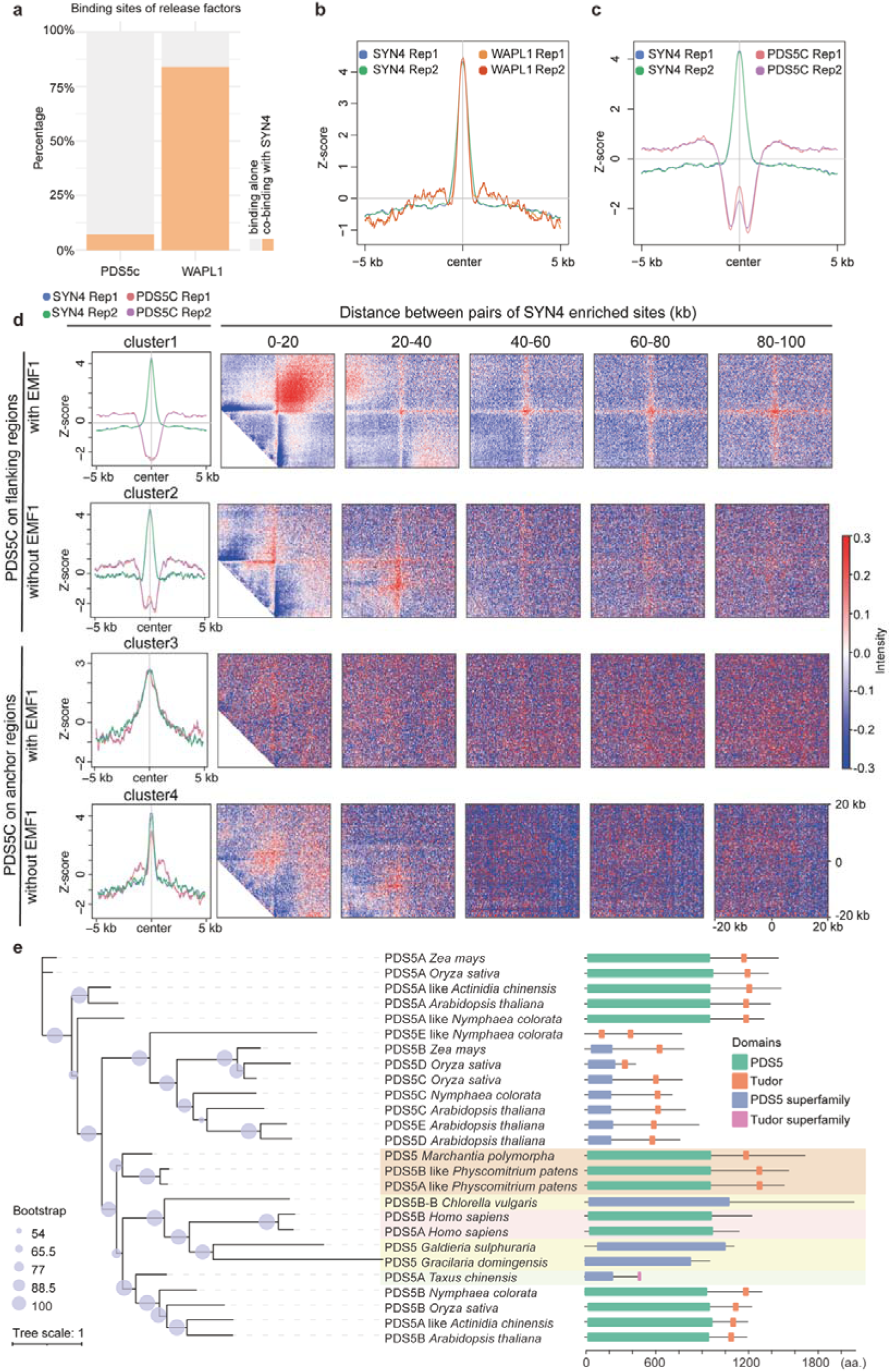
Distinct binding patterns of WAPL1 and PDS5C regulate noncanonical loops. a, Overlap ratios between WAPL1 and SYN4 binding sites and PDS5C and SYN4 binding sites. **b** and **c**, Metaplots showed ChIP-seq enrichment profiles for WAPL1 **(b)** and PDS5C **(c)** centered on SYN4 binding regions. **d,** APA analysis of four anchor categories in WT (1-kb resolution; 20-kb flanking regions): Anchors with EMF1 binding + PDS5C at flanking regions; Anchors lacking EMF1 + PDS5C at flanking regions; Anchors co-bound by EMF1 and PDS5C; Anchors with PDS5C binding + EMF1 absence. **e,** Evolutionary analysis of PDS5 proteins: left - phylogenetic tree of PDS5 orthologs from algae (yellow), bryophytes (orange), gymnosperms (green), angiosperms (white), and human (control, pink). Bootstrap support values scale with circle size. Right - conserved domain architecture.

Previous studies indicated that PDS5C specifically binds to H3K4me1^35^, raising the question of whether H3K4me1 distribution determines PDS5C’s characteristic boundary flanking binding pattern. To investigate this issue, we performed H3K4me1 ChIP-seq in the WT and *pds5a/b/c* mutants (Extended Data Fig.6a). Results showed H3K4me1 localized outside of SYN4 binding boundaries and was enriched in the regions flanking these boundaries (Extended Data Fig.6b). Given that mutation of H3K4me1-binding proteins (PDS5s) abolishes noncanonical loops, we further compared H3K4me1 levels between the WT and *pds5a/b/c* mutants to investigate whether H3K4me1 also regulates these loops. The H3K4me1 levels remained nearly unchanged in mutants relative to WT (Extended Data Fig.6c-f), demonstrating that noncanonical loop loss in *pds5a/b/c* results exclusively from protein defects rather than altered histone modification.

Evolutionary analyses identified two key plant-specific differences in cohesin associated factors that contribute to the presence of noncanonical loops in plants. First, previous studies revealed that plant WAPL proteins lack the critical FGF motif required for interaction with PDS5^34^, explaining their non-overlapping binding patterns. Second, *Arabidopsis* PDS5C’s distinct binding specificity is conferred by its Tudor domain-dependent recognition of H3K4me1^35^. To determine when plant PDS5s acquired this domain, we conducted a phylogenetic analysis spanning algal species, bryophytes, gymnosperms, and angiosperms (monocots/dicots), with human PDS5s as controls (Fig.4e). The Tudor domain in PDS5s has been present since the emergence of land plants, whereas the orthologs in algae lack this crucial domain. This result implies the widespread presence of noncanonical loops across land plants.

### Impact of noncanonical loops on chromatin architecture

In mammals, both TADs and fountain structures consist of via cohesin-dependent loops constrained within boundaries^11,44–46^. Given the unique presence of noncanonical loops in plants, we investigated their contribution to higher-order chromatin organization. TAD structures are hardly observed in WT plants and are largely abolished in the *syn4* mutant (Extended Data Fig.7a). As previously described, noncanonical loops persist and lengthen in *wapl1/2* mutant. Those elongated loops are also detectable in the heatmap analysis. However, lengthening loops do not enhance the TAD definition (Extended Data Fig.7a). Strikingly, the *pds5a/b/c* mutants exhibited a complete loss of noncanonical loops, accompanied by the appearance of sharper TAD structures (Fig.3c and Extended Data7a). When *PDS5s* are mutated, noncanonical loops are not constrained at boundary-flanking regions and extend toward subsequent boundaries. Consequently, noncanonical loops disappear and enlarged TADs form, using subsequent SYN4-binding sites as boundaries (Fig.5e and Extended Data Fig.7b). This reorganization demonstrates that noncanonical loop formation suppresses the development of pronounced TAD architecture in plants.

**Fig. 5:**
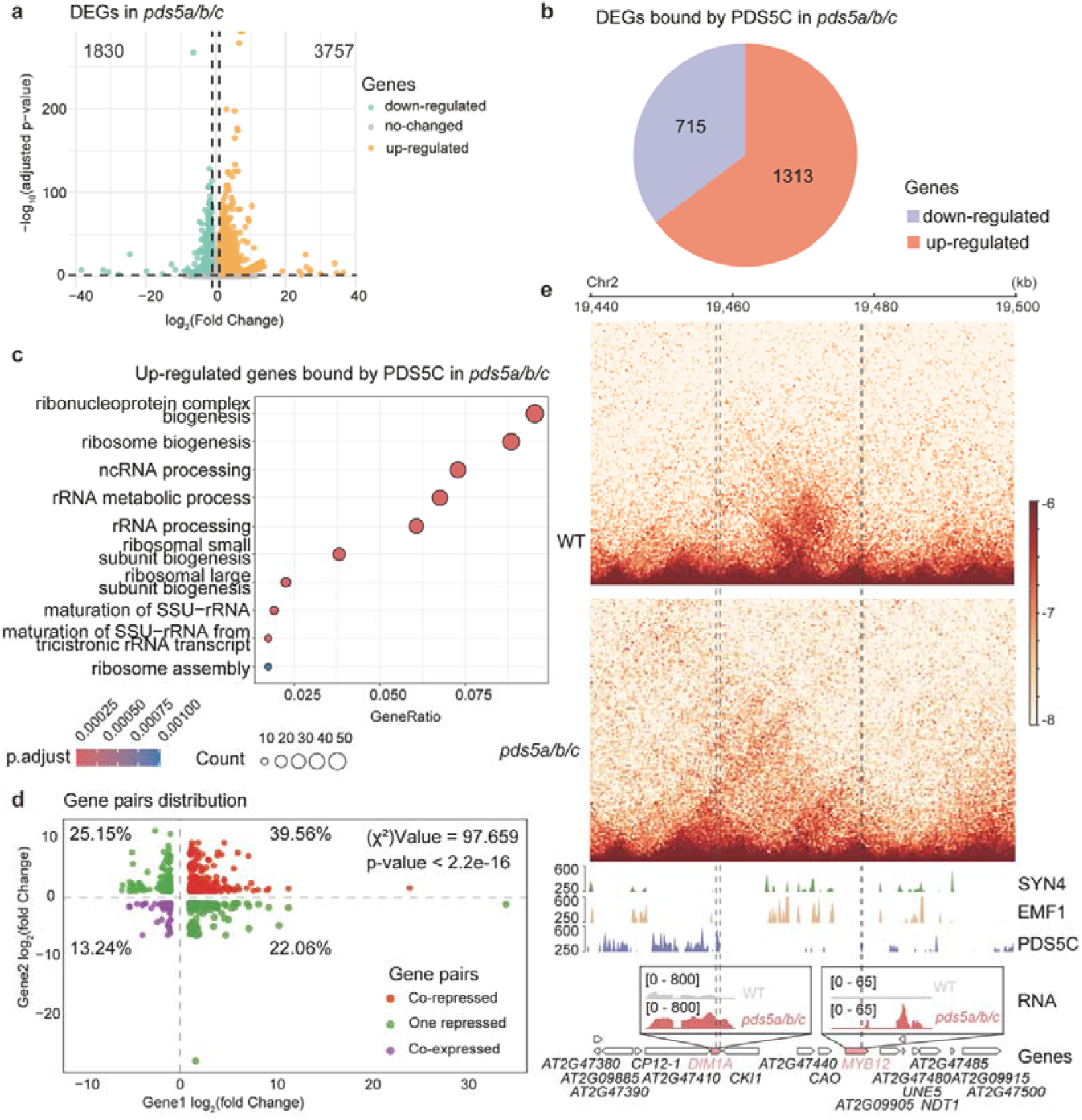
Noncanonical loops co-repress target gene pairs. a, Volcano plot of DEGs (q-value < 0.05 and |log_2_ fold change| > 1) in *pds5a/b/c* mutants compared to WT. **b,** PDS5C-targeted up-regulated and down-regulated genes in *pds5a/b/c* mutants are shown. **c,** GO enrichment analysis of PDS5C-targeted up-regulated genes in *pds5a/b/c*mutants. Dot size represents gene count, and color gradient indicates adjusted p-value (p.adjust). **d,** Scatterplot of log_2_ (fold change of *pds5a/b/c* and WT) for gene pairs anchored at noncanonical loops. **e,** Exemplary co-repressed gene pair regulated by noncanonical loops. Top: Micro-C heatmaps (200 bp resolution) showed noncanonical loop in WT (upper) and its absence in *pds5a/b/c* (lower). PDS5C enrichment occurs upstream and downstream 20-kb flanking regions of paired SYN4 anchors. SYN4 and EMF1 binding sites are annotated. Bottom: Transcript levels of target genes in WT versus *pds5a/b/c* mutant.

While mammalian fountains result from cohesin extrusion starting at fountain centers and stopping at boundaries (Extended Data Fig.8a)^45^, quantitative comparison of fountain strength across WT, *syn4*, *wapl1/2*, and *pds5a/b/c* mutants showed crucial differences: fountains completely disappear in *syn4*, confirming their dependence on cohesin, but are attenuated yet persistent in both *wapl1/2* and *pds5a/b/c* mutants (Extended Data Fig.8b). This result demonstrates that plant fountains are composite structures formed by both canonical and noncanonical loops. Unlike mammals, plant fountains involve the dynamic cycling of cohesin from the fountain center to WAPL-bound boundaries and PDS5-bound flanking regions (Extended Data Fig.8a).

### Noncanonical loops function to co-repress target gene pairs

After elucidating the mechanistic basis of noncanonical loop formation and its impact on higher-order chromatin architecture, we investigated their biological importance in plant development and identified their target genes. Differential expression analysis in *pds5a/b/c* revealed significantly more upregulated genes (n=3757) than downregulated genes (n=1830) (Fig.5a). PDS5C-bound differentially expressed genes (DEGs) showed a similar regulatory asymmetry, suggesting that PDS5s predominantly mediate transcriptional repression (Fig.5b). Gene Ontology (GO) enrichment analysis of upregulated targets showed strong associations with ribosome biogenesis processes (Fig.5c). To further examine the repressive function of noncanonical loops, we analyzed the transcription levels of noncanonical loop associated genes in *pds5a/b/c* mutants. First, we identified SYN4 peak anchors upstream and downstream separated by an intra-distance less than 20 kb. Then, we mapped PDS5C peaks located within the 20 kb upstream and downstream flanking regions of these anchors, respectively, thereby defining noncanonical loop anchors corresponding to the APA plot (cluster 1 in Fig.4d). These PDS5C peaks were annotated to genes, generating gene pairs associated with noncanonical loops (Supplementary Table 1). Analysis of these DEG pairs in *pds5a/b/c* mutants revealed that approximately 40% (389 genes) were co-upregulated, indicating that noncanonical loops function to co-repress anchor gene transcription in the WT (Fig.5d). Nearly all of these co-repressed genes (387/389) remained unchanged in the *wapl1/2* mutant (Extended Data Fig.9), in which noncanonical loops remained intact. This finding confirms that co-repression of these gene pairs requires noncanonical loops. For example, the gene pair *AT2G47420* (*ADENOSINE DIMETHYL TRANSFERASE 1A*, *DIM1A*) and *AT2G47460* (*MYB DOMAIN PROTEIN 12*, *MYB12*) is depicted in the Micro-C heatmaps for the WT (Fig.5e). DIM1A is reported as the only known 18S rRNA demethylase dedicated to ribosome biogenesis^47^. MYB12 functions as a transcription factor that regulates flavonoid accumulation in *Arabidopsis*^48^. This gene pair was anchored by a noncanonical loop in WT, with PDS5C binding to their gene bodies and detectable paired SYN4 signals between them; this contact was lost in *pds5a/b/c* (Fig.5e). The transcriptional levels of both genes were upregulated in *pds5a/b/c* mutant. Taken together, these findings demonstrate that noncanonical loops function to co-repress target gene pairs.

## Discussion

Several studies have shown that plants have distinct 3D chromatin organization compared to mammals^38–40,49^. However, the mechanism underlying this difference remains unclear. Our data reveal that *Arabidopsis* employs a distinct regulatory mechanism for cohesin-dependent loop formation. Most notably, unlike in mammals, boundaries are not impenetrable barriers in plants, permitting cohesin to extrude loops across boundaries until encountering PDS5 proteins at boundary-flanking regions (Fig.6a). The coexistence of canonical loops and noncanonical loops alters other cohesin-regulated 3D chromatin structures (such as TADs and fountains) in plants versus mammals, contributing to their unique chromatin organization. In *pds5a/b/c* mutants, which lack noncanonical loops, triangular TAD structures become sharper and fountain structures are weakened, resembling mammalian chromatin organization more closely (Fig.6b).

**Fig. 6:**
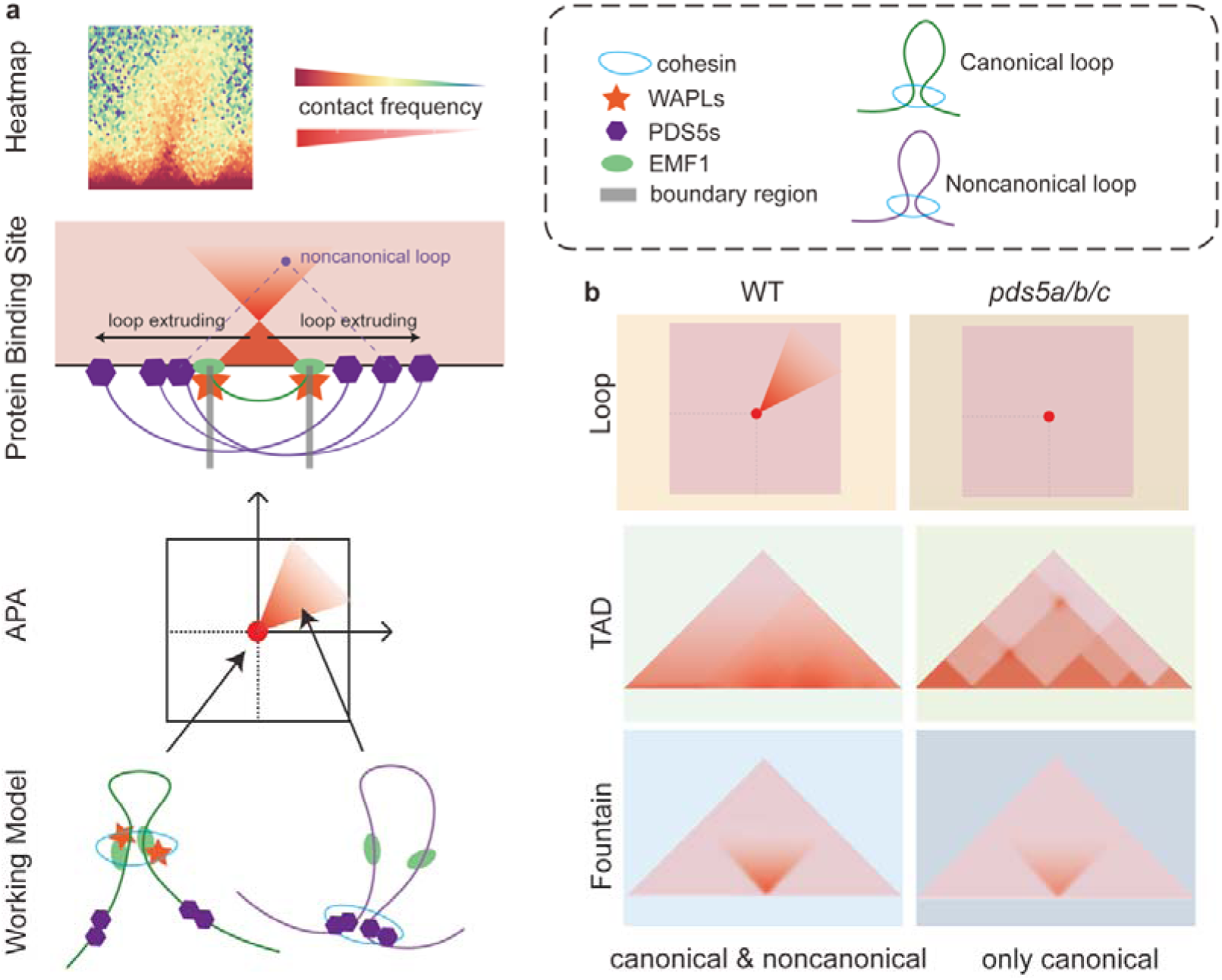
Plant-specific factors regulate noncanonical loops to shape distinct 3D genome architecture in *Arabidopsis.* a, The mechanism of forming canonical and noncanonical loops in *Arabidopsis*. The top heatmap shows an example for canonical and noncanonical loop in *Arabidopsis*. The schematic diagram shows the regulatory factors with their binding sites and how they function to shape the 3D chromatin structures in *Arabidopsis*. **b,** The schematic diagram shows the difference in loops, TADs and fountains between WT plant and *pds5a/b/c* mutant.

The dynamics of cohesin on chromatin are characterized by continuous cycles of loading, sliding, and release^11^. The aforementioned proteins synergistically regulate these cohesin dynamics^11^. Although cohesin’s loop extrusion function is conserved, evolutionary differences in CTCF and cohesin-associated factors lead to a distinct loop regulatory mechanism in plants. In mammals, CTCF serves as a boundary factor that halts cohesin at loop anchors through cohesin-CTCF interactions^42,50^. Moreover, single-nucleus Hi-C studies reveal that boundary positions vary across individual cells but statistically cluster at CTCF/cohesin co-binding sites^51,52^. This dynamic process of loop extrusion by cohesin is confined between boundaries^11,53^. In *Arabidopsis*, no CTCF homologs exist. Instead, the plant-specific boundary-binding protein EMF1 interacts with the cohesin complex. However, these interactions likely have lower affinity than mammalian cohesin-CTCF interactions, resulting in diminished anchor confinement capacity and permitting increased cohesin mobility across anchors. Furthermore, in mammals, WAPL and PDS5 proteins interact and cooperate with CTCF to inhibit extrusion of longer cohesin-dependent loops at boundaries^12,15^. In contrast, in *Arabidopsis*, WAPL and PDS5 homologs exhibit substantial structural divergence. WAPL proteins retain the C-terminal WAPL domain and cohesin-release function^34^. Consequently, they still release cohesin at boundary regions and are essential for forming canonical loops. Critically, the N-terminal region of plant WAPL proteins lacks the FGF motif present in mammalian homologs that mediates interaction with PDS5 proteins; this absence prevents interaction with PDS5 proteins^34^. Meanwhile, *Arabidopsis* PDS5 proteins possess a Tudor domain, enabling direct binding to boundary-flanking regions enriched with H3K4me1^35^, specifically rather than to boundaries themselves. This likely leads to inefficient cohesin release at boundaries, increasing the possibility of loop extrusion across boundaries. PDS5 proteins subsequently restrict elongation of these noncanonical loops, and their binding sites constitute the anchors of those loops.

In conclusion, our study demonstrates that noncanonical loops define the distinctive chromatin architecture of plants. This organizational mechanism relies on two plant-specific adaptations: the boundary protein EMF1 and PDS5s containing H3K4me1-binding Tudor domains. Noncanonical loops co-repress specific gene pairs and play critical roles in regulating plant development.

## Methods

### Plant materials and growth conditions

The mutants and WT plants used in this study were derived from the Columbia-0 (Col-0) background. Several mutants have been previously described: *syn4* (SALK_130085), *emf1-2*^54^, *wapl1-1* (SALK_076791), *wapl2* (SALK_127445), *pds5a* (SALK_114556), *pds5b* (SALK_092843), and *pds5c* (SALK_013481) (Supplementary Table 2). The *wapl1/2* and *pds5a/b/c* mutants were generated by cross. For all experiments, seeds were sown on Murashige and Skoog (MS) plates containing 1% sucrose and 0.7% agar, stratified at 4°C for 2 days, and then grown for 10 days under long-day conditions (8 h dark/16 h light) at 22°C. Whole seedlings were used for experiments.

### Micro-C

Micro-C libraries were constructed following established protocols^41^. Approximately 2 g of plant tissue were ground into a fine powder in liquid nitrogen and incubated with Nuclei Isolation Buffer lysis solution at 4°C for 15 min. The homogenate was filtered through a 40-µm cell strainer; nuclei were cross-linked with 3% formaldehyde (FA) for 15 min at 4°C, followed by quenching with 0.375 M glycine for 5 min. FA-fixed nuclei were subjected to additional cross-linking with freshly prepared 3 mM ethylene glycol bis (succinimidyl succinate) (EGS) for 60 min at 4°C. The dual cross-linked nuclei (FA + EGS) were quenched with 0.4 M glycine for 15 min at 4°C and washed twice with phosphate-buffered saline containing 0.05% bovine serum albumin. Chromatin was fragmented using micrococcal nuclease (MNase) based on pre-titrated digestion tests and incubated for 10 min at 37°C. The reaction was stopped by adding 1.5 mM egtazic acid (EGTA) at 65°C for 10 min. DNA 3’ ends were dephosphorylated and 5’ ends were phosphorylated using 20 units of T4 polynucleotide kinase (NEB, #M0201L) at 37°C for 15 min. DNA overhangs were filled in using 40 units of DNA Polymerase I, Large (Klenow) Fragment (NEB, #M0210L) with biotin-dCTP and biotin-dATP (Invitrogen) at 25°C for 45 min. The reaction was terminated by incubation with 0.03 M ethylenediaminetetraacetic acid (EDTA) at 65°C for 20 min. The chromatin was collected and subjected to proximity ligation using 50 units of T4 DNA Ligase (NEB, #M0202L) at room temperature for 3 h. After ligation, unligated biotin-labeled DNA ends were removed using 200 units of exonuclease III (NEB, #M0206S) at 37°C for 5 min. Cross-links were reversed by treatment with proteinase K at 65°C for 2 h. The ligated DNA was then extracted using the Qiagen DNeasy Plant Mini Kit (Qiagen, #69106), in accordance with the manufacturer’s instructions. DNA fragments of 250–400 bp were selected and purified using 0.7X + 0.3X AMPure XP beads. The fragments underwent blunt-end repair, polyadenylation, and adaptor ligation using the VAHTS Universal Plus DNA Library Prep Kit for MGI (Vazyme, #NDM617). Subsequently, streptavidin C1 beads (Invitrogen, #65001) were used for pull-down, followed by polymerase chain reaction (PCR) amplification with the following conditions: 95°C for 3 min; 10–12 cycles of 98°C for 20 s, 60°C for 15 s, and 72°C for 30 s; 72°C for 5 min; hold at 4°C. Amplified products were purified with 0.8X AMPure XP beads. The final Micro-C libraries were quantified and sequenced on the MGI-seq platform (BGI, China).

### RNA-seq

Total RNA was extracted using the E.Z.N.A. Plant RNA Kit (Omega, R6827-01). Libraries were prepared using VAHTS® mRNA Capture Beads (Vazyme, N401) and the VAHTS® Universal V8 RNA-Seq Library Prep Kit for MGI (Vazyme, NRM605). Sequencing was performed on the MGI DNBSEQ T7 PE150 platform. Each sample included three biological replicates.

### ChIP-seq

For WAPL1, a 7,325 bp genomic DNA fragment, including 2,000 bp upstream of the WAPL1 start codon and lacking the stop codon, was PCR-amplified using specific primers (Supplementary Table 2) and recombined into the 1300-FLAG vector. ChIP-seq experiments were performed as previously described^55^. Briefly, 10-day-old seedlings were fixed with 1% formaldehyde, flash-frozen in liquid nitrogen, and nuclei were isolated using nuclear isolation buffer (NIB: 50 mM HEPES pH 8.0, 5% sucrose, 5 mM MgCl_2_, 5 mM NaCl, 40% glycerol, 0.25% Triton X-100, 0.1 mM phenylmethylsulfonyl fluoride, 0.1% β-mercaptoethanol), then filtered through double-layered Miracloth. Chromatin was fragmented by sonication in TE-SDS buffer (10 mM Tris-HCl pH 7.4, 1 mM EDTA, 0.25% sodium dodecyl sulfate [SDS]) to produce 200–500 bp fragments. Immunoprecipitations were performed using specific antibodies: H3K4me1 (Abcam ab8895) and FLAG (Millipore F7425) in IP buffer (80 mM Tris-HCl pH 7.4, 230 mM NaCl, 1.7% NP-40, 0.17% DOC). Antibody–chromatin complexes were captured with rProtein A Sepharose Fast Flow beads, then subjected to five stringent washes. After overnight de-crosslinking at 65°C, DNA was purified by phenol–chloroform extraction and ethanol precipitation.

Two biological replicates per immunoprecipitation were processed for library construction using the VAHTS® Universal Pro DNA Library Prep Kit (Vazyme NDM608). Library preparation included end repair, A-tailing, and ligation of Illumina-compatible adapters, followed by 14 cycles of PCR amplification. Size selection (200–600 bp) was performed using VAHTS™ DNA Clean Beads (N411). Final libraries were sequenced on the MGI DNBSEQ T7 PE150 platform, generating 2 × 150 bp paired-end reads.

### RNA-seq bioinformatics analysis

Raw reads were processed using fastp^56^, then mapped with gaps to the reference genome (TAIR10, https://www.arabidopsis.org) using HISAT2^57^. SAMtools was used for filtering, indexing, and file format conversion^58^. Reads were annotated using StringTie^59^ based on the genome reference (Araport11, https://www.arabidopsis.org). Library correlations were calculated using Pearson correlation. Differentially expressed genes were identified using the R package DESeq2^60^. GO enrichment analysis was conducted using clusterProfiler^61^. Transcript abundance was quantified via FPKM normalization in deepTools^62^. Representative genomic loci were visualized with the Integrative Genomics Viewer (IGV)^63^.

### Phylogenic analysis

PDS5 protein sequences of various species were downloaded from NCBI (https://www.ncbi.nlm.nih.gov/protein/) and UniProt (https://www.uniprot.org/), then aligned using MAFFT^64^. The phylogenic tree was generated via IQTREE^65^. The parameter -m MFP was applied to identify the best-fit model^66^, and JTT+F+I+G4 was selected to reconstruct the tree with 1,000 ultrafast bootstrap replicates^67^. The phylogenic tree was visualized using iTOL (https://itol.embl.de). Conserved domains were predicted and visualized using the NCBI CD-Search Tool^68^ and Chiplot (https://www.chiplot.online/).

### ChIP-seq bioinformatics analysis

Sequencing data were preprocessed using fastp^56^ to remove adapter sequences and low-quality reads. Clean reads were aligned to the *Arabidopsis* reference genome (https://www.arabidopsis.org) using Bowtie^69^, allowing a maximum of two nucleotide mismatches. PCR duplicates were identified and marked using MarkDuplicates in Picard (http://broadinstitute.github.io/picard/). SAMtools was utilized to filter uniquely mapped reads for downstream analyses^58^. Peaks were identified using MACS2^70^.

Normalized genome coverage profiles were generated via deepTools bamCoverage with a bin size of 10 bp, RPKM normalization, and effective genome size compensation.^62^ Representative genomic loci were visualized using IGV^63^. To assess reproducibility among ChIP-seq replicates, multiBigwigSummary and plotCorrelation tools were used with a bin size of 1,000. Signal intensities in meta-plots were calculated as log_2_(IP/input) RPKM ratios and visualized using SeqPlots^71^. The R package ChIPpeakAnno^72^ was used for peak annotation.

### Hi-C bioinformatics analysis

Low-quality reads were filtered with fastp^56^, and the remaining reads were mapped to the reference genome (TAIR10, https://www.arabidopsis.org) using Bowtie2^73^. Valid pairs were identified using HiC-Pro with default parameters^74^. A multiple fixed-resolution contact matrix was generated and balanced using cooler^75^. The contact matrix was visualized with CoolBox^76^.

### Micro-C bioinformatics analysis

After low-quality data had been processed using fastp^56^, we referred to published articles^41^ and the pipeline described at https://micro-c.readthedocs.io to complete data processing. Alignment of libraries to the reference genome was performed using BWA-MEM^77^. Subsequently, pairtools^78^ was used to identify and filter ligation events, generating valid pairs based on the following criteria: mapq > 40, gap < 30 bp, and no duplication. The resulting .pairs file was converted to .cool format using pairix (https://github.com/4dn-dcic/pairix). A multiple fixed-resolution matrix was generated and balanced using cooler^75^. For comparisons of contact maps among samples, sequencing depths were normalized to approximately 400 million valid pairs per sample for downstream analysis. Contact probability at different genomic distances was quantified using IDE, calculated with HiCExplorer^79^. Contact heatmaps were visualized using CoolBox^76^ and HiGlass (https://github.com/higlass/higlass-python).

### Aggregate peak analysis (APA)

APA was performed using the cooltools framework^80^, which allowed customized analyses. Briefly, all pairwise interactions between target anchors within a specified distance range were identified and extracted as observed interactions. Simultaneously, interactions between randomly selected site pairs within the same range were calculated to establish a random background baseline. Observed interactions were normalized against this background to generate APA results. The APA heatmap was produced at a 1-kb resolution with a 20-kb flanking region around each target site. In the heatmap, red indicates stronger interactions between target sites, whereas blue represents a higher frequency of interactions within the random background. The identification of fountain structures was described in the previous study^36^.

## Data availability

All sequencing data generated in this study have been deposited in the Genome Sequence Archive (GSA) at the National Genomics Data Center (NGDC, PRJCA033743) and are publicly accessible at https://ngdc.cncb.ac.cn/gsa. The SYN4, PDS5C, and EMF1 ChIP-Seq raw data were obtained from previously published studies^31,32,35^. The RNA-seq raw data for *pds5a/b/c* mutants were acquired from other researchers^33^. The Micro-C raw data for the WT were also obtained from a published study^41^.

## Acknowledgments

This study was conducted at the Peking University High-Performance Computing Platform, and calculations were performed on CLS-HPC. This work was supported by the Biological Breeding-National Science and Technology Major Project (2023ZD04073, Yue Zhou); grant 32370612 (Yue Zhou) from the National Natural Science Foundation of China; grant JCTD-2022-06 (Yue Zhou and Yuannian Jiao) from the CAS Youth Interdisciplinary Team; and startup funds from the State Key Laboratory of Gene Function and Modulation Research, the School of Advanced Agricultural Sciences, and the Peking-Tsinghua Center for Life Sciences at Peking University (Yue Zhou). Additionally, this study was supported by grant PID2022-142997NB-I00 from MCIN/AEI /10.13039/501100011033 / FEDER, UE (Myriam Calonje).

## Author information

These authors contributed equally: Dingyue Wang, Suxin Xiao, Lingxiao Luo, Jiayue Shu, Guangmei Tian.

## Ethics declarations

The authors declare no competing interests.

## Extended data

Extended Data Figures 1-9.

## Supplementary information

Supplementary Tables

Supplementary Tables 1-2.

